# Molecular characterization of a complex of Apoptosis Inducing Factor 1 (AIFM1) with cytochrome c oxidase of the mitochondrial respiratory chain

**DOI:** 10.1101/2021.03.31.437858

**Authors:** Johannes F. Hevler, Riccardo Zenezeni Chiozzi, Alfredo Cabrera-Orefice, Ulrich Brandt, Susanne Arnold, Albert J.R. Heck

## Abstract

Combining mass spectrometry based chemical cross-linking and complexome profiling, we analyzed the interactome of heart mitochondria. We focused on complexes of oxidative phosphorylation and found that dimeric apoptosis inducing factor 1 (AIFM1) forms a defined complex with ~10% of monomeric cytochrome *c* oxidase (COX), but hardly interacts with respiratory chain supercomplexes. Multiple AIFM1 inter-crosslinks engaging six different COX subunits provided structural restraints to build a detailed atomic model of the COX-AIFM1_2_ complex. Application of two complementary proteomic approaches thus provided unexpected insight into the macromolecular organization of the mitochondrial complexome. Our structural model excludes direct electron transfer between AIFM1 and COX. Notably however, the binding site of cytochrome *c* remains accessible allowing formation of a ternary complex. The discovery of the previously overlooked COX-AIFM1_2_ complex and clues provided by the structural model hint at a role of AIFM1 in OXPHOS biogenesis and in programmed cell death.

## Introduction

Mitochondria are considered the powerhouse of aerobic eukaryotic cells, as they contain the major pathways of oxidative energy metabolism and produce the bulk of ATP by oxidative phosphorylation (OXPHOS) necessary for cellular homeostasis. Only at the end of the last century it became evident that mitochondria also are key players in apoptosis and that this process is tightly linked to OXPHOS components (Saraste, 1999). Apoptosis Inducing Factor (AIFM1) was one of the proteins found to be released from the mitochondrial intermembrane space during programmed cell death and to have the capacity to induce chromatin condensation and DNA fragmentation in a caspase-independent fashion (Susin et al., 1999). A mutation found in AIFM1 has been associated with Cowchock syndrome (OMIM 310490) (Rinaldi et al., 2012). Early on, it was also reported that ablation of AIFM1 leads to OXPHOS deficiency (Vahsen et al., 2004), in line with findings that AIFM1 mutations cause combined oxidative phosphorylation deficiency 6, a severe mitochondrial encephalomyopathy (OMIM 300816; (Ghezzi et al., 2010). More recently, it has been proposed that AIFM1 is involved in the disulfide relay of the mitochondrial intermembrane space by serving as import receptor of CHCHD4/MIA40 (Hangen et al., 2015; Meyer et al., 2015; Petrungaro et al., 2015). However, the specific mechanisms and molecular interactions by which these different functions of AIFM1 are connected in health and disease are not well resolved. For example, AIFM1 deficiency affects OXPHOS predominantly by lowering the amount of respiratory chain complex I (Vahsen et al., 2004). Other components were found to be affected in a tissue specific manner. In AIFM1 deficient patients (Ghezzi et al., 2010), ablation of AIFM1 in skeletal and heart muscle affected cytochrome c oxidase (COX) in addition to complex I, whereas in liver a deficiency of complexes I and V was observed (Joza et al., 2005; Pospisilik et al., 2007). In the present study, we explored the molecular interactions of AIFM1 with the multiprotein complexes of the OXPHOS system in heart mitochondria using our recently established complementary experimental approach (Hevler et al., 2021) that combines cross-linking mass-spectrometry (XL-MS) and complexome profiling (Figure 1).

**Figure 1.**
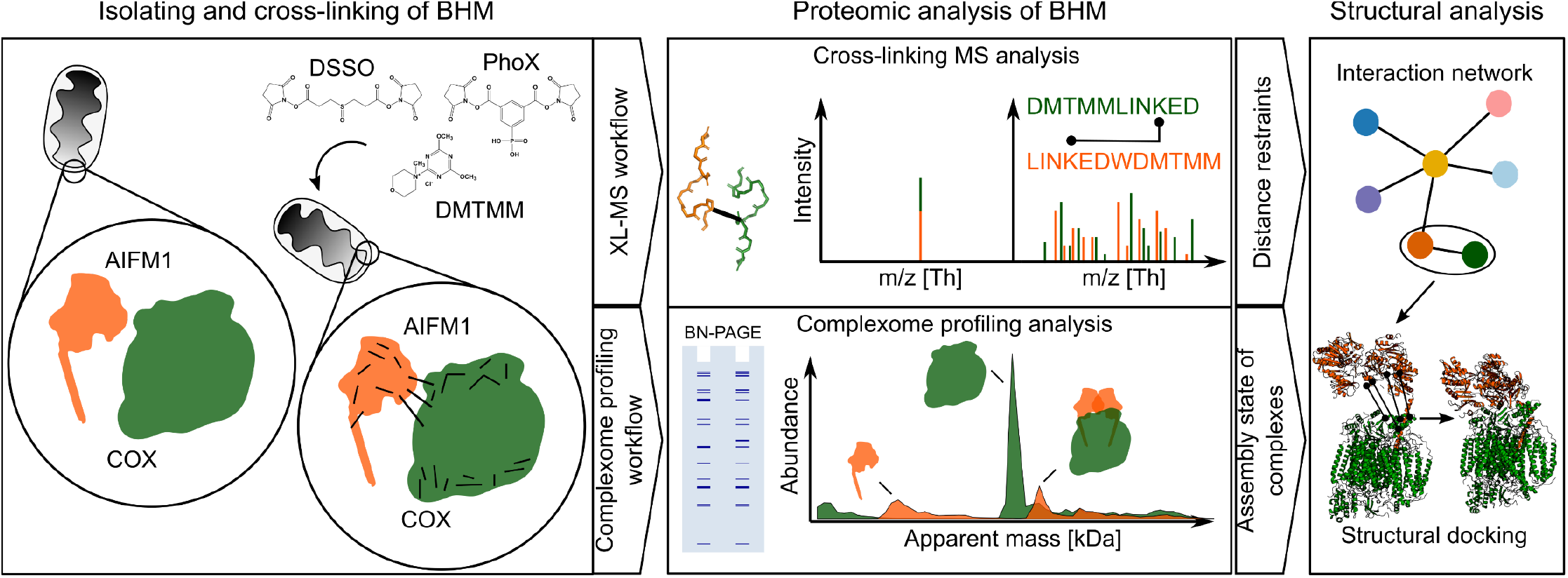
Two-tier experimental strategy for the analysis of proteome-wide protein-protein interactions in bovine heart mitochondria. Mitochondria membranes were cross-linked with either of the three cross-linking reagents DSSO, PhoX or DMTMM. Subsequently, samples were analyzed by cross-linking mass spectrometry (XL-MS) and complexome profiling. Identified cross-linked peptides were used to generate protein-protein interaction networks. Protein interactions and structural models of AIFM1 with COX were then computationally modelled using the distance restraints from XL-MS data together with the assembly state and stoichiometry information obtained by complexome profiling.

To increase the depth and confidence of the study, bovine heart mitochondrial membranes (BHM) were treated with three different chemical cross-linkers: DSSO (Kao et al., 2011), PhoX (Steigenberger et al., 2019) and DMTMM (Leitner et al., 2014). While DSSO and PhoX predominantly generate lysine-lysine residue cross-links, DMTMM acts as a condensation reagent of acidic side chains of aspartic or glutamic acids with lysine side chains, resulting in the formation of a stable bond between those residues. We found that a significant fraction of AIFM1 in its dimeric form is specifically bound to monomeric cytochrome *c* oxidase, an interaction that has been overlooked so far. By using the identified cross-links as structural restraints, we generated a structural model of dimeric AIFM1 docked to monomeric COX.

## Results and Discussion

We analyzed the organization and interaction landscape of protein complexes in bovine heart mitochondria by combining cross-linking mass spectrometry XL-MS and complexome profiling (Heide et al., 2012; Hevler et al., 2021) thereby adding new information on native state multiprotein complexes of interest, building upon previous work exploring the interactome of mitochondria from different organisms, tissues and cells by XL-MS (Chavez et al., 2020; Linden et al., 2020; Liu et al., 2018; Ryl et al., 2020; Schweppe et al., 2017).

### Exploring mitochondrial complexes by combined cross-linking and complexome profiling

To increase the depth of the protein-interaction map of BHM we applied multiple cross-linkers (DSSO, PhoX and DMTMM). Covering 215 proteins listed in MitoCarta 3.0 (Rath et al., 2020), we obtained a total of 4413 unique cross-links (3261 intra- and 1152 inter-protein cross-links; Figure S1a-b, Appendix Table 1). In accordance with previously published studies (Fursch et al., 2020; Gonzalez-Lozano et al., 2020), the abundance of detected cross-linked proteins was higher than the median of all identified proteins in the BHM sample (8.8 vs. 6.9 log_10_ iBAQ; Figure S1b). Reflecting the large number and high abundance of membrane integral multiprotein complexes and the very high protein density, especially within the inner mitochondrial membrane, ~75% of the cross-links identified involved membrane proteins (Figure S1c). For the same reasons and corroborating previous studies using mouse and human mitochondria (Liu et al., 2018; Rath et al., 2020; Ryl et al., 2020; Schweppe et al., 2017) the largest number of cross-links reflected interactions between the many subunits of OXPHOS complexes and their association to supercomplexes of respiratory chain complexes I, III and IV (1431 out of 4131 cross-links), also called respirasomes (Schägger and Pfeiffer, 2000) (Figure S1d).

Complexome profiling analysis of untreated (i.e. non-cross-linked) BHM yielded very similar results as those obtained previously with rat heart mitochondria using the same approach (Heide et al., 2012), showing very similar migration pattern of the OXPHOS complexes and respirasomes (Figure S1e, Appendix Table 2). When the samples were cross-linked with PhoX and DMTMM before subjecting them to complexome profiling, the overall abundance of detected proteins was not affected substantially. However, it was evident from the migration profiles of OXPHOS complexes that cross-linking to some extent prevented dissociation of complex V (CV) dimers and other fragile higher order respiratory supercomplexes during native electrophoresis (Figure S1e). Importantly, in most cases the apparent molecular masses of the bulk of the OXPHOS complexes were not markedly affected by cross-linking. A notable exception was complex III (CIII) in the DMTMM treated sample, where it migrated not predominantly at the apparent mass of its obligatory dimer at ~500 kDa as in all other conditions, but showed at ~650 kDa and multiple peaks at higher masses. The shift of CIII-dimer to higher masses suggested that, possibly through the large hydrophilic domains of its two core subunits, this OXPHOS complex cross-linked to a much larger extent to other mitochondrial proteins, than the others. The ~800 kDa peak corresponds to a previously described supercomplex between one complex III dimer and one complex IV (COX) monomer (Chatzispyrou et al., 2018). The latter was also found in untreated and PhoX cross-linked samples, but was much more pronounced after cross-linking with DMTMM. The peaks at ~1100 kDa and ~1300 kDa can be interpreted as dimers of complex III dimers without and with one monomer of COX, respectively.

Taken together, these results establish that classical XL-MS analysis alone and in combination with complexome profiling delivered consistent results. Separating native complexes prior to mass spectrometric analysis provided additional key information on their apparent molecular masses and multimeric state. Cross-linking them beforehand allowed for more reliable detection of more fragile assemblies that otherwise partially or completely dissociate during solubilization and/or electrophoresis.

### A specific complex between AIFM1 dimers and COX revealed by XL-MS

When we performed an in-depth analysis of all detected cross-links involving OXPHOS complexes in addition to engaging their canonical components themselves, one specific protein stood out: In all our XL-MS datasets combined, AIFM1 had inter-links with no less than six subunits of COX with 82% of them involving COX6B1 and COX6C (Figure 2a, Appendix Table 3). Cross-links with COX subunits accounted for 86% of the total inter cross-links with AIFM1. Adenylate kinase 2 (AK2) and adenine nucleotide carrier isoform 1 (SLC25A4) were the only other two proteins featuring multiple inter-protein cross-links with AIFM1.

**Figure 2.**
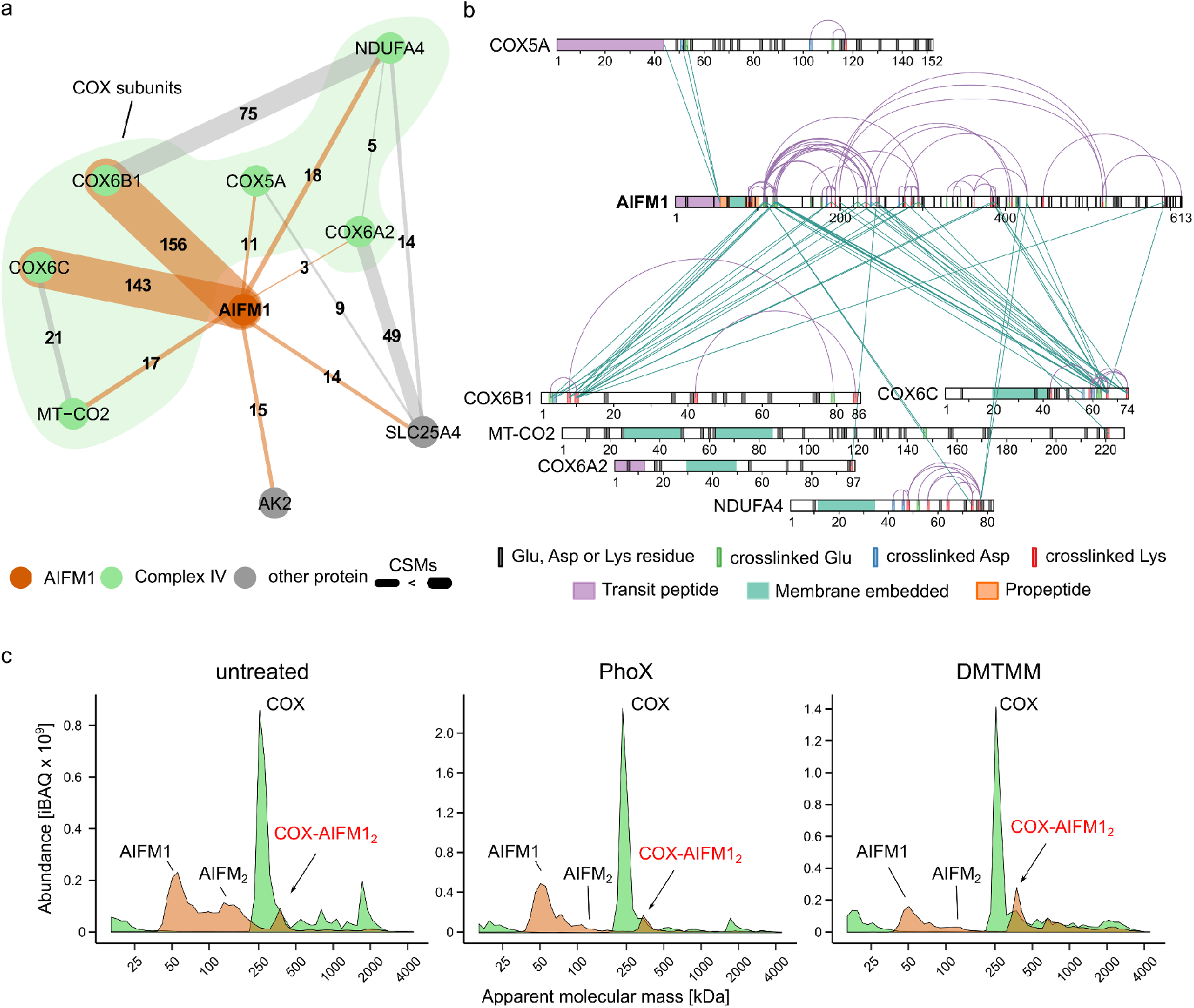
Dimeric AIFM1 forms a defined complex with monomeric COX. **a)** Interaction network of AIFM1 in cross-linked BHM. Numbers and thickness of lines indicate the cumulative evidence (CSMs) for each interaction. Orange lines indicate cross-links involving AIFM1, while cross-links between AIFM1 interactors are presented as gray lines. **b)** Xi-net plot of the COX-AIFM1 interaction. Purple colored links indicate intra-links. Green colored links indicate inter-links. Respective sequence and cross-link features are indicated accordingly. **c)** Migration profiles of AIFM1 (orange) and averaged COX (green) from untreated and cross-linked (PhoX or DMTMM) mitochondria separated by BN-PAGE (4-16%). Peaks are annotated based on the apparent molecular mass of AIFM1 (~62 kDa) and monomeric COX (~ 220 kDa). In all samples, peaks corresponding to monomeric AIFM1 and COX as well as a peak corresponding to a COX-AIFM1_2_ complex are observed. Treatment with DMTMM seems to prevent dissociation of the COX-AIFM1_2_ complex.

The association of AIFM1 with this OXPHOS complex is remarkable in particular, since COX from bovine heart is undoubtedly the longest and best studied version of cytochrome c oxidase (Arnold, 2012). Therefore, we interrogated an earlier cross-linking dataset of mouse heart mitochondria for this interaction (Liu et al., 2018). Corroborating our findings, the majority of AIFM1 cross-links identified in this study engaged three different COX subunits, with COX6C being the most prominent by far (Figure S2a). Of note, Liu and coworkers detected multiple cross-links between AIFM1 and AK2 as well in mouse heart mitochondria. Moreover, charting large affinity purifications-mass spectrometry (AP-MS) depositories, we found that they contained multiple instances of COX subunits interacting with AIFM1 (Antonicka et al., 2020; Hein et al., 2015; Huttlin et al., 2017). Yet, buried in datasets generated by large-scale analyses of the mitochondrial interactome, these indications for AIFM1 binding to COX seem to go unnoticed so far.

Detailed evaluation of the observed cross-links between AIFM1 and COX (Figure 2b, Figure S2b) revealed that they were predominantly within the pyridine nucleotide-disulfide oxidoreductase domain (Pfam: PF07992; res 136-460) of AIFM1 comprising one FAD- and one NADH-binding domain. Suggesting that AIFM1 had not been cleaved to its truncated pro-apoptogenic form (Sevrioukova, 2011), additional intra- and inter-protein cross-links were observed at the N-terminal end of the pro-peptide (res 55-101) of AIFM1 that is predicted to cross the inner mitochondrial membrane (IMM) reaching to the matrix side. Notably, these cross-links were the only ones to the matrix facing subunit COX5A, while all other cross-links engaged domains of COX subunits facing the intermembrane space (IMS).

Our three independent cross-linking analyses strongly suggested that AIFM1 and COX formed a specific complex, but provided no information on the multimeric state of the interaction partners and how much of this unexpected complex was present in bovine heart mitochondria. Therefore, we applied complexome profiling to analyze complexes containing AIFM1 and COX using, the same samples as in the XL-MS analysis (Figure 2c; Appendix Table 2). In all cases, COX was predominantly present as a monomer (~220 kDa) and a prominent fraction of AIFM1 was found to migrate at an apparent mass consistent with its monomeric state (62 kDa)..

Substantial amounts of AIFM1 dimers were only observed in untreated BHM indicating that they may be destabilized by the cross-linking protocol. This was possibly due to partial oxidation of NADH known to be required for AIFM1 dimerization (Hangen et al., 2015). Importantly however, a significant amount of AIFM1 consistently showed up as a peak at an apparent mass of ~350 kDa in untreated mitochondria as well as after cross-linking with PhoX or DMTMM. This peak coincided with a shoulder next to the prominent peak at ~220 kDa of monomeric COX in all samples analyzed, thus suggesting the presence of a ~350 kDa complex containing a dimer of AIFM1 (~124 kDa) bound to monomeric COX (~220 kDa). Notably, a shoulder on the higher mass side of the COX monomer can also be observed in complexome profiling data of human cells published earlier, but its significance was not evident at the time (Guerrero-Castillo et al., 2017). Label free quantification revealed that hardly any of the other respiratory chain complexes were present in this segment of the migration profiles. In contrast, the amounts of the COX monomer and AIFM1 dimer were comparable at ~350 kDa suggesting a stoichiometric association, reflecting the observations for gels with increased resolution (high range BN-gel (3-10%)) (Figure S2c). At the same time, no AIFM1 co-migrated with the bulk of monomeric COX at ~220 kDa (Figure S2d). Notably, only small amounts of AIFM1 were detected at ~1,850 kDa, the predicted mass of supercomplex S1 (I_1_III_2_IV_1_). This was mostly observed in the DMTMM treated sample that exerted many more inter-protein cross-links in the high mass range overall (Figure S1a, d). It can be concluded that AIFM1_2_ was in our samples bound almost exclusively to monomeric COX and, if any, very little could be found associated with supercomplexes. Consistent with its higher cross-linking efficiency, the fraction of COX engaged in the complex with AIFM1_2_ was somewhat higher with DMTMM than in the untreated and PhoX cross-linked samples. In fact, in untreated samples the amount of COX-AIFM1_2_ complex was variable to some extent. This suggested that it tended to dissociate during solubilization and native electrophoresis. Such behavior has been observed previously for several less tightly associated subunits of OXPHOS complexes (Abdrakhmanova et al., 2005; Balsa et al., 2012; Hirst et al., 2003). Overall, we could estimate that about 10% of monomeric COX was engaged in a fairly stable stoichiometric complex with AIFM1 dimers (Figure S2d).

In summary, combination of cross-linking and complexome profiling data provided compelling evidence for the presence of a defined COX-AIFM1_2_ complex in bovine heart mitochondria. The interaction interface was defined as involving residues of the neighboring COX6C, COX6B1, NDUFA4, COX6A2 and MT-CO2 contacting the pyridine nucleotide-disulfide oxidoreductase domain of AIFM1, and residues of COX5A interacting with the matrix facing N-terminal region of its pro-peptide.

### Creation of a cross-link guided structural model of the COX-AIFM12 complex

Next, we aimed at building a structural model for the COX-AIFM1_2_ complex guided by the distance restraints obtained by cross-linking, also including those involving the N-terminal sequence of AIFM1 comprising its pro-peptide sequence (res 55-101). We first derived a *de novo* model of this so far structurally unresolved region using trRosetta (Yang et al., 2020) to complement a homology model of the bovine AIFM1 dimer that we derived from the human structure (PDB: 4BUR; (Ferreira et al., 2014)) using Robetta (Kim et al., 2004). The structural model obtained for the N-terminal domain of AIFM1 agreed well with secondary structure predictions and featured three alpha-helices (res 67-88; 94-98; 105-112) of which the first is predicted to be a transmembrane segment (Figure S3a). We then used restraints derived from our cross-linking data to dock this model of the bovine AIFM1 dimer to the 1.8 Å structure (PDB: 1V54) of COX (Tsukihara et al., 2003). Unfortunately, this COX structure does not contain the more loosely attached NDUFA4 subunit. Therefore, we used Robetta (Kim et al., 2004) to complement it with a homology model derived from the human NDUFA4 structure (Zong et al., 2018)(PDB: 5Z62 chain N).

Mapping of the cross-links (intra-links) onto the structural models of AIFM1_2_ and COX revealed that the majority of cross-links were below 30 Å, with a combined mean distance of 19.1 Å for DSSO/PhoX and 20.8 Å for DMTMM cross-links (Figure 3a, Figure S3b). The mean distance for DSSO and PhoX cross-links was well within the theoretical maximal range of ~30 Å and ~25 Å, respectively. DMTMM cross-links averaged somewhat above the theoretical maximum of ~15 Å, in line with previous observations (Leitner et al., 2014). Note, that 8 cross-links for AIFM1 and 10 cross-links for COX were not included in these calculations, because they involved intra-links from AIFM1 (res 128-613) to its *de novo* modelled N-terminus (res 55-124) or regions not resolved in the structural models (AIFM1 res 517-550; COX4l1 res 23-25; Figure S3b).

**Figure 3.**
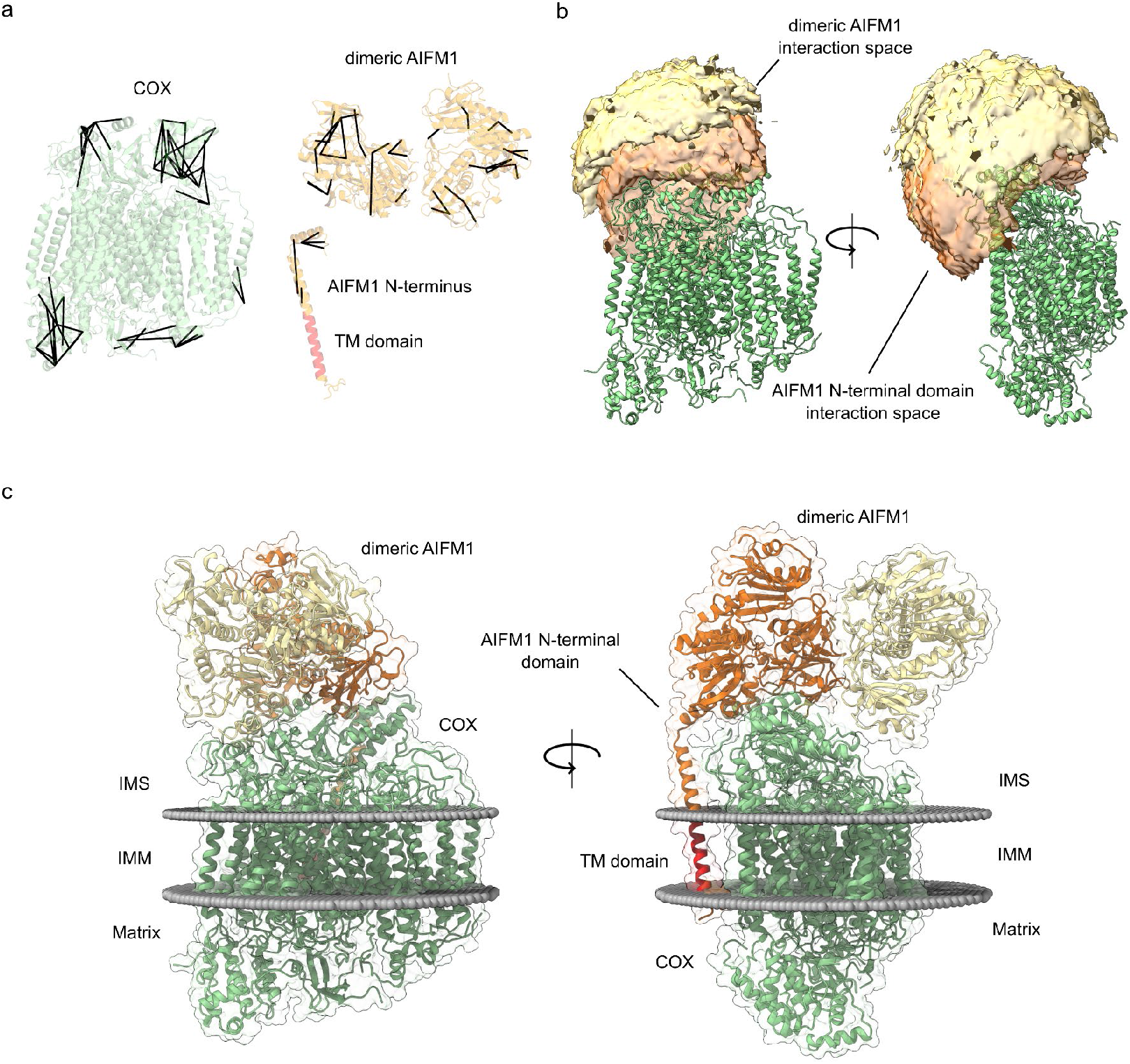
Cross-link derived structural model of the COX-AIFM1_2_ complex. **a)** Detected intra-links mapped onto the structural models of COX, AIFM1 dimer as well as the de-novo modelled N-terminal domain of AIFM1. The COX structure (green) was based on a previously resolved structure (PDB 1V54) supplemented with NDUFA4 (structurally aligned based on PDB: 5Z62). Dimeric AIFM1 (orange) was modelled based on the previously resolved human homologue (PDB 4BUR, res 128-516, 551-613). The N-terminal domain of AIFM1 (orange) (res 55-124) was generated using trRosetta. The predicted transmembrane (TM) domain is highlighted in red. **b)** Visualization of the cross-link driven accessible interaction space models for a COX-AIFM1_2_ complex. COX is represented in green, while the bright orange volume represents the center-of-mass position of the AIFM1 dimer, and the dark orange volume represents the center-of-mass position of the model of the AIFM1-N-terminus (res 55-124). The cross-linking data are consistent with the interaction space available for docking dimeric AIFM1 and the N-terminal region of one AIFM1 protomer to monomeric COX. **c)** Cross-link derived structural model of the COX-AIFM1_2_ complex. COX is represented in green, and AIFM1 protomers (res 128-516; 551-613 with and without N-terminal region (res 55-127) are represented in orange and yellow respectively. The transmembrane residues (67-85) of the N-terminus of the interacting AIFM1 moiety is highlighted in red. Membrane boundaries of the IMM are sketched as gray spheres. The final complex consists of monomeric COX, dimeric AIFM1 (res 128-516, 551-613) and the N-terminal region of one AIFM1 protomer (res 55-127)

Based on the solvent accessibility and distance restraints obtained from both structures, accessible interaction interfaces between COX and the AIFM1 dimer as well as COX and the N-terminal region of AIFM1 were calculated using DisVis (van Zundert and Bonvin, 2015). While this analysis suggested that the AIFM1 dimer attaches to the intermembrane space side of COX, the predicted interaction space for N-terminal region of one AIFM1 protomer covers the transmembrane domain at the matrix side of COX making contacts to the COX6B1, COX6C MT-CO2, NDUFA4 and COX5A subunits (Figure 3b).

Scoring the interface models using the restraints imposed by the cross-linking data, suggests that the COX-AIFM1_2_ interface is mostly occupied by just one AIFM1 protomer. In agreement with this notion, cross-links suggested that only one N-terminal region, not both of the AIFM1 dimer interacted directly with COX. Therefore, the final modelling of the COX-AIFM1_2_ complex was performed by docking just one AIFM1 protomer and one N-terminal region of AIFM1 to the COX monomer. Cross-link driven docking using Haddock (van Zundert et al., 2016) resulted in seven distinct clusters. The model best satisfying both, the cross-linking data and presenting the highest Haddock score was chosen to represent the COX-AIFM1_2_ complex (Appendix Table 4). Overall, 46 of 59 unique cross-links were used eventually to dock AIFM1_2_ and one of its N-terminal domains with monomeric COX. In the final structural model of the COX-AIFM1_2_ complex, the AIFM1 dimer “sits” on COX facing the intermembrane space side and makes contact through one AIFM1 protomer covering parts of COX6B1, COX6C1, MT-CO2 and NDUFA4 (Figure 3c). The second AIFM1 protomer points away from COX making just one very limited contact to COX through its C-terminal loop. At the opposite side of COX, the N-terminal region of the interacting AIFM1 protomer makes contact with COX6B1 and transmembrane helices of MT-CO2 and NDUFA4 (Figure 3c).

It should be noted that after docking with Haddock, the *de novo* structural model covers the N-terminal region of AIFM1 only up to residue 124 creating a structurally undefined stretch of three amino acids up to residue 128, the first amino acid contained in the homology model for the main part of AIFM1. Therefore, residues 121 to 131 were re-modelled using the “Model Loops” interface in Chimera (Pettersen et al., 2004; Yang et al., 2012), thereby connecting the N-terminal domain to the main part of AIFM1. Cross-links (46) used for structural modelling as well as all observed cross-links (59) for COX-AIFM1_2_ were in good agreement with the predicted structure (Figure S3c). Cross-links used for structural docking with a distance >35 Å (13 cross-links, maximum distance 48.7 Å) mostly included cross-links from the N-terminal domain (res 55-124) to a flexible segment of COX6C (6 cross-links) as well as to the FAD-binding domain of AIFM1 (4 cross-links), supporting an even closer localization of the intermembrane N-terminal domain of AIFM1 to COX (Figure S3d).

Next, we performed and interaction interface analysis of the docking model that predicted three distinct interfaces between COX and AIFM1 (Figure 4a, Appendix Table 5). The first, extensive interface is defined by the N-terminal residues of AIFM1, which interacts with neighboring residues of the COX subunits MT-CO2, NDUFA4 and COX6B1. Secondly, residues of the pyridine nucleotide-disulfide oxidoreductase domain of AIFM1 comprising the NADH- and FAD-binding domains intimately interact with MT-CO2, COX6B1 and COX6C. The third, rather small interaction interface is defined by residues of the C-terminal region of the second AIFM1 protomer and residues of MT-CO2, COX4l1 and COX7B (Figure 4b). The interface between the hydrophilic parts of the N-terminal region of AIFM1 facing the intermembrane space is mostly driven by contacts to COX6B1 and NDUFA4, whereas its transmembrane domain predominantly interacts with one of the transmembrane segments of MT-CO2. Notably, the very N-terminal residues of AIFM1 facing the matrix side reside within a ~25 Å distance from residues 44-54 of COX5A, consistent with the observed cross-links to this COX subunit (Figure 2b).

**Figure 4.**
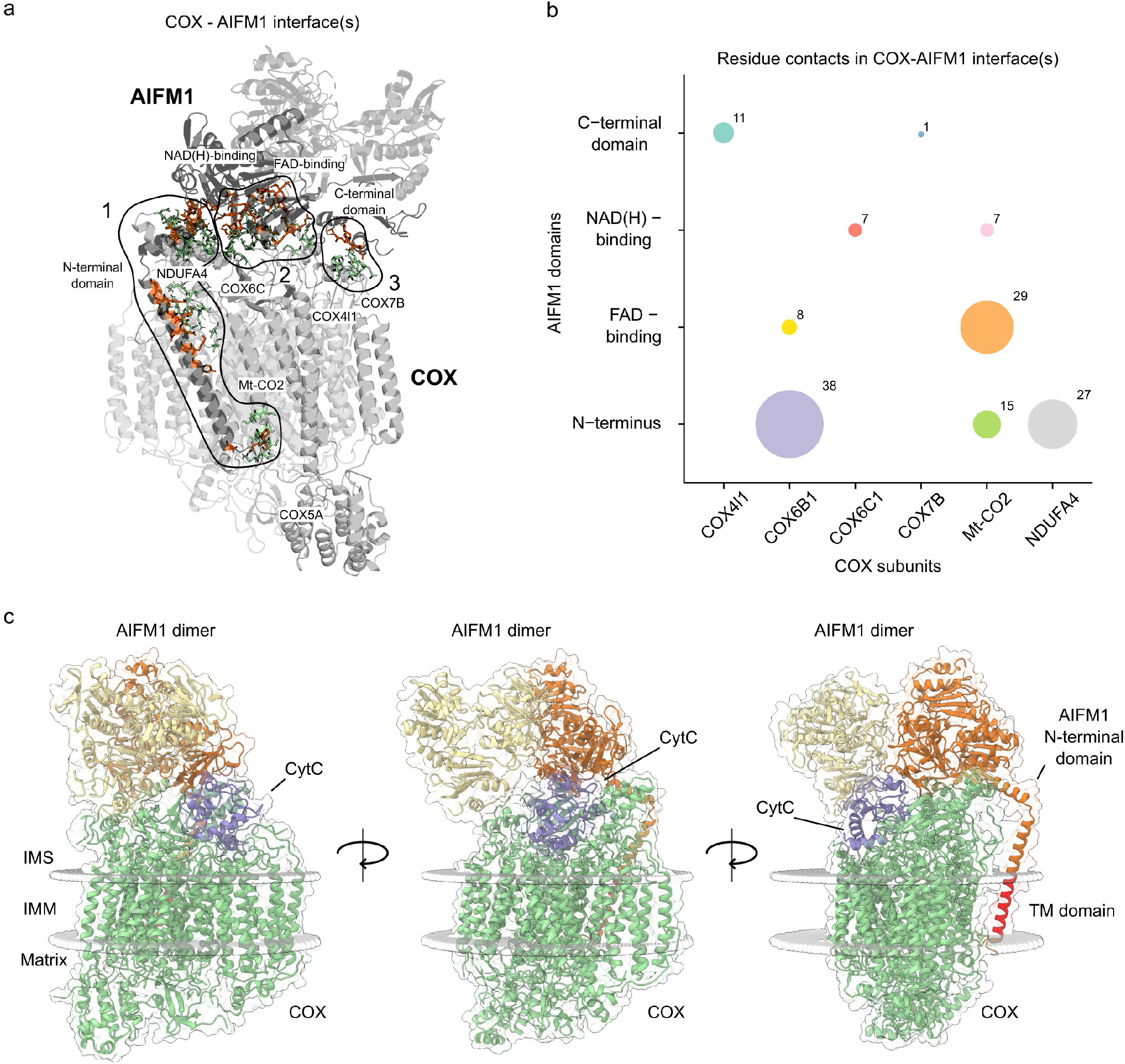
Deciphering interaction interfaces in the COX-AIFM1_2_ structural model. **a)** Three distinct interfaces between COX subunits and respective AIFM1 protomers were found. Subunits (COX) and protein domains (AIFM1) with residues in respective interfaces are colored in gray. Active COX residues are shown as green colored sticks and active AIFM1 residues as orange colored sticks. **b)** Analysis of the number of residue contacts between respective COX subunits and AIFM1 domains. Colored circles indicate residue contacts between single subunits (COX) and domains (AIFM1) with the size of each circle corresponding to the number of residue-residue interactions. **c)** COX-AIFM1_2_ complex with cytochrome *c* (purple) bound to its COX binding site. The structural model presented here was merged with a previously published model of cytochrome *c* docked to COX from bovine heart (Sato et al., 2016). COX subunits are colored green, while AIFM1 protomers are colored orange and yellow. The transmembrane (TM) domain of the N-terminal domain of AIFM1 is highlighted in red. Boundaries of the IMM are indicated as gray spheres.

### Potential functional implications of a COX-AIFM12 complex

While the predominant consequence of AIFM1 deficiency is impaired complex I assembly (Vahsen et al., 2004), additional COX deficiency has been reported in skeletal muscle and heart (Joza et al., 2005; Pospisilik et al., 2007), as well as in *Drosophila melanogaster* (Joza et al., 2008) and *Caenorhabditis elegans* (Troulinaki et al., 2018). Conversely, AIFM1 expression was found to be significantly increased along with several COX assembly factors in human COX-negative muscle fibers (Murgia et al., 2019). However, it seems unlikely that the COX-AIFM1_2_ complex described here contributes to assembly or stabilization of COX, because it accounted only for 10% or less of the total amount of this OXPHOS complex. Moreover, no apparent co-migration between AIFM1 and any of the individual COX subunits or sub-assemblies at apparent masses lower than ~350 kDa was observed, suggesting that association of AIFM1 occurred only with fully assembled COX. Conditional involvement of AIFM1 in the maturation of COX assembly factors that are substrates of the disulfide relay of the intermembrane space (Hangen et al., 2015; Meyer et al., 2015; Petrungaro et al., 2015) appears as a more likely explanation for the link between AIFM1 and COX deficiency in some tissues.

It has been reported that AIFM1 is a member of the NDH-2 family of proteins (Kerscher, 2000) and thus exhibits NADH:ubiquinone oxidoreductase activity (Elguindy and Nakamaru-Ogiso, 2015). Nevertheless, we examined the possibility of direct electron transfer between the FAD and CuA of COX within the COX-AIFM1_2_ complex. The minimal distance between the isoalloxazine moieties of the FADs and the Cu_A_ center was ~50 Å and ~55 Å (Figure S4), which is more than three-times larger than the 14 Å considered as the maximum distance for efficient electron tunneling in a protein matrix (Moser et al., 2010).

It would be conceivable that this distance is bridged by cytochrome *c* serving as an electron shuttle between AIFM1 and COX. Therefore, we explored whether cytochrome *c* could still bind to its substrate binding site in the COX-AIFM1_2_ complex. Merging our structural model with a previously obtained model of cytochrome *c* bound to COX from bovine heart (Sato et al., 2016) suggested that the AIFM1 dimer does not hamper cytochrome c from binding to COX (Figure 4c). However, distances of ~45 Å and ~47 Å between the heme moiety of cytochrome *c* and the isoalloxazine rings of FAD in both AIFM1 protomers excluded direct electron transfer also in the presence of the additional heme. Yet, a substrate channeling mechanism could still be imaginable implying movement of cytochrome *c* forth and back between AIFM1 and COX without leaving the complex. A crevice in the second AIFM1 protomer facing COX could potentially reduce the distance between the redox centers indeed to about 14 Å. However, this would require cytochrome *c* to turn within the pocket formed by AIFM1_2_ and COX in order to bring its heme as close as possible to the isoalloxazine ring. Thus, while such a substrate channeling mechanism cannot be excluded, it does not seem very likely. Moreover, oxidizing NADH would transiently destabilize dimerization of AIFM1 (Hangen et al., 2015) and thus the entire complex, arguing further against any oxidoreductase activity of the COX-AIFM1_2_ complex. If COX-AIFM1_2_ is not a catalytic complex, it is still tempting to speculate that a ternary interaction of COX, AIFM1_2_ and cytochrome *c* could play a role in mitochondrial pro-apoptotic mechanisms. Apart from directly promoting programmed cell death (Cande et al., 2002; Cheung et al., 2006; Ghezzi et al., 2010), AIFM1 could play an indirect role in apoptosis by modulating release of cytochrome *c* (Saraste, 1999) through its binding to the COX-AIFM1_2_ complex. For this, it is important to note that cytochrome *c* makes direct contact to the first AIFM1 protomer in the ternary complex (Figure 4c). Providing further support for this hypothesis, cross-links between AIFM1 and cytochrome *c* were previously reported in intact mouse heart mitochondria (Liu et al., 2018). It is known that binding of cytochrome *c* to COX is strongly reduced at higher ionic strength (Sinjorgo et al., 1986), which could explain, why we did not observe cross-links between cytochrome *c* (CytC) and the COX-AIFM1_2_ complex. Since the N-terminal pro-peptide with its transmembrane helix provides a significant portion of the AIFM1/COX interface, the complex is expected to destabilize upon cleavage of AIFM1 thereby activating its pro-apoptotic function. In addition to cleaved AIFM1, any previously bound cytochrome *c* would be released concomitantly further promoting apoptosis, potentially providing a synergistic boost to the cell death program already underway.

## Conclusions

We show that ~10% of monomeric COX in bovine heart mitochondria are engaged in a defined complex with dimeric AIFM1. Using structural restraints provided by cross-linking data, available high-resolution structures and structural modeling, we could derive a model of the COX-AIFM1_2_ complex with and without bound cytochrome *c*. Combining chemical cross-linking and complexome profiling provided useful complementary information and represents proof-of-concept for our experimental approach demonstrating that it can be used to define and characterize multiprotein assemblies in detail that may have been overlooked by other means.

While our structural model excludes direct electron transfer between AIFM1 and COX, it provides clues on potential functional implications of the formation of the COX-AIFM1_2_ complex including a possible involvement in promoting apoptosis. The structural insights into this unexpected mitochondrial complex will stimulate and guide further studies on the role of AIFM1 in OXPHOS biogenesis and apoptosis.

## Data availability

All cross linking mass spectrometry related data, the structural docking (Haddock results), and the presented COX-AIFM1/ COX-AIFM1-CytC structural model described in this work have been deposited to the ProteomeXchange partner PRIDE database) and assigned the identifier PXD025102 (Vizcaino et al., 2016). Complexome profiling datasets have been deposited in the CEDAR database (van Strien et al., 2021).

## Acknowledgments

All authors acknowledge support from the Netherlands Organization for Scientific Research (NWO) funding the Netherlands Proteomics Centre through the X-omics Road Map program (Project 184.034.019) and the TOP project 714.017.004, the Netherlands Organization for Health Research and Development (ZonMW TOP 91217009), the EU Horizon 2020 program Epic-XS (Project 823839), and the German Research Foundation (DFG) through the Collaborative Research Center 1218 (Project 269925409).

## Author contributions

JFH, UB, SA and AJRH designed the study. JFH and RCZ performed XL-MS experiments. JFH performed XL-MS analysis and structural modeling. ACO performed the complexome profiling experiments. ACO, SA and UB analyzed the complexome profiling datasets. ACO and SA provided the bovine mitochondrial samples. JFH, SA and AJRH wrote the original draft, with all authors carefully revising and editing the manuscript before submission. UB, SA and AJRH acquired funding and resources. AJRH supervised the project.

## Materials and Methods

### Isolation and purification of bovine heart mitochondria (BHM)

Mitochondrial membranes from bovine heart were isolated and preserved as described in (Hevler et al., 2021). In order to increase the purity of the preparation and for Tris-buffer removal, frozen crude mitochondria (4 × 15 ml aliquots; 60 mg protein/ml) were thawed on ice, diluted (1:4) with ice-cold SEH buffer (250 mM sucrose, 1 mM EDTA, 20 mM HEPES, pH 7.4 adjusted with NaOH) and centrifuged at 1,000 × g (10 min; 4°C). The supernatants were recovered and centrifuged at 40,000 × g (20 min; 4°C) and each resulting pellet was suspended in 2 ml SEH buffer. Afterwards, mitochondria were loaded onto a two-layer sucrose gradient (1 M sucrose, 20 mM HEPES, pH 7.4 /1.5 M sucrose, 20 mM HEPES, pH 7.4) and centrifuged at 60,000 × g (20 min; 4°C). The pure mitochondrial fractions accumulated at the interphase were carefully recovered and pooled into one tube. After resuspension in 20 ml ice-cold SEH buffer, pure mitochondria were centrifuged at 10,000 × g (20 min; 4°C) and finally suspended in 5 ml ice-cold SEH buffer supplemented with protease inhibitor cocktail (SIGMAFAST™). Protein concentration was determined by the DC protein assay (Bio-Rad) and aliquots of pure mitochondria were shock-frozen in liquid nitrogen and stored at −80°C until use.

### Cross-linking of BHM sample with DSSO, PhoX and DMTMM

Purified bovine heart mitochondrial membranes were buffer exchanged into cross-linking buffer (10 mM HEPES pH 7.8, 1 mM EDTA, 1 mM EGTA, 10 mM NaCl, 150 mM KCl, protease inhibitor). After optimization of the cross-link reaction, ~ 2 mg of BHM were either incubated with DSSO (0.5 mM freshly re-suspended in anhydrous DMSO; Thermo Fisher Scientific), PhoX (1 mM freshly re-suspended in anhydrous DMSO; made in-house) or DMTMM (10 mM freshly re-suspended in cross-linking buffer; Sigma-Aldrich) in 2 ml of cross-linking buffer at room temperature (RT). The cross-link reaction was quenched after 30 min by the addition of 50 mM Tris (1 M Tris buffer, pH 8.5) for additional 30 min at RT.

### Sample preparation for XL-MS analysis of cross-linked BHM

Cross-linked mitochondria were solubilized with Digitonin (9 g/g protein) for 30-60 min on ice. Proteins were denatured and purified as described previously (Leung et al., 2021). Briefly, denatured proteins were re-suspended and digested overnight (ON) at 37°C with Lys-C followed by Trypsin. The final peptide mixtures were desalted with solid-phase extraction C18 columns (Sep-Pak, Waters). Samples cross-linked with DSSO and DMTMM were fractionated with an Agilent 1200 HPLC pump system (Agilent) coupled to an strong cation exchange separation column (Luna SCX 5 μm – 100 Å particles, 50 × 2mm, Phenomenex), resulting in 24 fractions. For PhoX crosslinking we used a Fe^3+^-IMAC column (Propac IMAC-10 4 × 50 mm column, Thermo Fisher scientific) connected to an Agilent HPLC. Lyophilized peptides were dissolved in buffer A (30% acetonitrile, 0.07% trifluoroacetic acid) and the pH was adjusted to a value of 2. PhoX cross-linked peptides were subsequently eluted with a gradient of elution buffer B (0.3% NH_4_OH) (Potel et al., 2018). The collected PhoX-enriched peptides were then dried down and further fractionated into 7 high-pH fractions as previously described (Ruprecht et al., 2017).

### XL-MS analysis and data analysis

The 24 SCX fractions of DSSO were injected in an Agilent 1290 Infinity UHPLC system (Agilent) on a 50-cm analytical column packed with C18 beads (Dr Maisch Reprosil C18, 3 μm) coupled online to an Orbitrap Fusion Lumos (Thermo Fisher Scientific). We used the following LC-MS/MS parameters: after 5 minutes of loading with 100% buffer A (water with 0.1% formic acid), peptides were eluted at 300 nL/min with a 97 minutes gradient from 4% to 39% of buffer B (80% Acetonitrile and 20% water with 0.1% formic acid). For MS acquisition we used a MS1 Orbitrap scan at 120,000 resolution from 310 to 1600, AGC target of 5e^5^ ions and maximum injection time of 50 ms. The ions with a charge from +3 to +8 were fragmented with CID (NCE of 30%) and analyzed with MS2 Orbitrap at 30,000 resolution, AGC target of 5e^4^ ions and maximum injection time of 54 ms for detection of DSSO signature peaks (difference in mass of 37.972 Da). The four ions with this specific difference were analyzed with a MS3 Ion Trap scans (AGC target of 2e^4^ ions, maximum injection time of 150 ms) for sequencing the individual peptides. For the fractions of DMTMM and PhoX, we used an Ultimate3000 (Thermo Fisher Scientific) and 50-cm analytical column packed with C18 beads (Dr Maisch Reprosil C18, 3 μm) heated at 45°C, connected to Orbitrap Fusion Lumos. For both experiment we used a gradient from 9% to 40%, but in case of DMTMM it was 90 minutes long while for PhoX 30 minutes. For both experiments, we used a MS1 Orbitrap scan at 120,000 resolution from 350 to 1400, AGC target of 1e^6^ ions and maximum injection time of 50 ms. The most abundant ions with a charge between +3 and +8 were fragmented in HCD (stepped NCE of 30±3%) and analyzed with MS2 Orbitrap scan at 30,000 resolution, AGC target of 1e^5^ ions, and maximum injection time of 120 ms. The DSSO fractions were analyzed with Proteome Discoverer software suite version 2.4.1.15 (Thermo Fisher Scientific) with the incorporated XlinkX node for analysis of cross-linked peptides as reported by Klykov et al. (Klykov et al., 2018). Data were searched against a FASTA file containing the ~4200 most abundant proteins, which were previously determined following a classical bottom-up workflow. Were applicable, mitochondrial target peptides were removed from respective protein sequences. For XlinkX search, we selected fully tryptic digestion with three maximum missed cleavages, 10 ppm error for MS1, 20 ppm for MS2 and 0.5 Da for MS3 in Ion Trap. For modifications, we used static Carbamidomethyl (C) and dynamic Oxidation (M). The cross-linked peptides were accepted with a minimum score of 40, minimum score difference of 4 and maximum FDR rate set to 5%. Both non-cleavable cross-linkers were analyzed with pLink2 (Chen et al., 2019) and the same FASTA used for DSSO. For PhoX, we manually added the cross-linker to the list (alpha/beta sites “[K”, linker composition C(8)H(3)O(5)P(1) mass of 209.971Da) and for both cross-linkers, the same settings as described for XlinkX (without the minimum score option) was used. Finally, cross-links were additionally filtered: only cross-links corresponding to protein-protein interactions that were reported for at least two cross-linkers and with at least two CSMs were kept for the final interaction analysis and structural modeling.

### Complexome profiling analysis

Aliquots of untreated and PhoX and DMTMM cross-linked mitochondrial membranes (see “Cross-linking of BHM sample with DSSO, PhoX and DMTMM” for details) were thawed on ice, solubilized with Digitonin (9 g/g protein) in 50 mM NaCl, 50 mM imidazole-HCl, 2 mM 6-aminohexanoic acid, 1 mM EDTA, pH 7 and kept on ice for 20 min. Samples were further centrifuged at 22,000 × g (20 min; 4°C) and the supernatants were transferred into clean tubes and supplemented with Coomassie blue loading dye as described in Wittig et al. (2006). For blue-native (BN)-PAGE, 100 μg protein of each sample were loaded onto 4-16% or 3-10% polyacrylamide gradient gels and separated as described previously (Wittig et al., 2006). After the electrophoretic run, the gel was fixed ON in 50% methanol, 10% acetic acid, 100 mM ammonium acetate followed by staining with 0.025% Coomassie blue G-250 (Serva G) in 10% acetic acid for 30 min, destained twice in 10% acetic acid (1 h each) and kept in deionized water ON. The next day, the gel was color-scanned using a flatbed Image Scanner III (GE, USA) to use it as a template for the cutting procedure.

Proteins were identified by LC-MS/MS after in-gel tryptic digestion following the protocol described in Heide et al. (2012) with some modifications. In short, each gel lane was cut into 60 even slices starting at the bottom of the gel. The slices were cubed and transferred into 96-well filter plates (Millipore®, MABVN1250) adapted manually to 96-well plates (MaxiSorp™ Nunc) as waste collectors. Gel pieces were incubated with 50% methanol, 50 mM ammonium hydrogen carbonate (AHC) under moderate shaking; the solution was refreshed until the blue dye was removed completely. Removal of excess solution was done by centrifugation (1,000 × g, 15 s). In the next step, gel pieces were reduced with 10 mM dithiothreitol in 50 mM AHC for 1 h. After removing excess solution, 30 mM chloroacetamide in 50 mM AHC was added to each well, incubated in the dark for 45 min and removed. A short incubation step with 50% methanol, 50 mM AHC was performed for gel pieces dehydration (~15 min). The latter solution was removed and gel pieces were dried for ~30 min at RT. Later, 20 μl of 5 ng μl^−1^ trypsin (sequencing grade, Promega®) in 50 mM AHC plus 1 mM CaCl_2_ were added to each well and incubated for 20 min at 4°C. Gel pieces were covered by adding 50 μl of 50 mM AHC followed by an ON incubation at 37°C for protein digestion. The next day, the peptide-containing supernatants were collected by centrifugation (1,000 × g, 30 s) into clean 96-well PCR plates (Axygen®). The gel pieces were finally incubated with 50 μl of 30% acetonitrile (ACN), 3% formic acid (FA) for ~30 min prior elution of the remaining peptides on the previous eluates by centrifugation. The peptides were dried in a SpeedVac Concentrator Plus (Eppendorf) for 2.5-3 hours, resuspended in 20 μl of 5% ACN, 0.5% FA and stored at −20 °C until MS analysis.

After thawing the frozen resuspended peptides and a 30 min gentle shaking, individual samples were loaded and separated by reverse phase liquid chromatography and analyzed by tandem mass spectrometry in a Q-Exactive Orbitrap Mass Spectrometer equipped with a nano-flow ultra-HPLC system (Easy nLC1000, Thermo Fisher Scientific). In brief, peptides were separated using 100 μm ID × 15 cm length PicoTip^™^ EMITTER columns (New Objective) filled with ReproSil-Pur C18-AQ reverse-phase beads (3 μm, 120Å) (Dr. Maisch GmbH, Germany) using linear gradients of 5%–35% ACN, 0.1% FA (30 min) at a flow rate of 300 nl min^−1^, followed by 35%-80% ACN, 0.1% FA (5 min) at 600 nl min^−1^ and a final column wash with 80% ACN (5 min) at 600 nl min^−1^. All settings for the mass spectrometer operation were the same as detailed in (Guerrero-Castillo et al., 2017).

MS raw data files from all individual slices were analyzed using MaxQuant (v1.5.0.25) against the *Bos taurus* proteome entries retrieved from Uniprot. The following settings were applied: Trypsin, as the protease, N-terminal acetylation and methionine oxidation as variable modifications; cysteine carbamidomethylation as fixed modification; two trypsin missed cleavages; matching between runs, 2 min matching time window; six residues as minimal peptide length; common contaminants included; I = L and the rest of parameters were kept as default. Individual protein abundances were determined by label-free quantification using the obtained intensity-based absolute quantification (iBAQ) values, which were corrected for protein loading and MS sensitivity variations using the sum of total iBAQ values from each sample. For each protein group entry, migration profiles were generated and normalized to the maximal abundance through all fractions. The migration patterns of the identified proteins were hierarchically clustered by an average linkage algorithm with centered Pearson correlation distance measures using Cluster 3.0 (de Hoon et al., 2004). The resulting complexome profiles consisting of a list of proteins arranged according to the similarity of their migration patterns in BN-PAGE were visualized as heat maps representing the normalized abundance in each gel slice by a three-color gradient (black/yellow/red) and processed in Microsoft Excel for analysis. The mass calibration for the BN gel was performed using the apparent molecular masses of either membrane or soluble bovine heart mitochondrial proteins. For membrane proteins: VDAC1 (30 kDa), complex II (123 kDa), complex IV (215 kDa), complex III (dimer, 485 kDa), complex V (700 kDa), complex I (1000 kDa), respiratory supercomplexes, I-IV (1215 kDa), I-III_2_ (S0, 1485 kDa), I-III_2_-IV (S1, 1700 kDa), I-III_2_-IV_2_ (S2, 1915 kDa) and complex V tetramer (2400 kDa). For Soluble proteins: ATP synthase subunit beta (51 kDa), citrate synthase (dimer, 98 kDa), ETFA/B (dimer, 122 kDa), enoyl-CoA hydratase (hexamer, 169 kDa), fumarase (tetramer, 200 kDa), Heat shock protein 60 (heptamer, 406 kDa), PCCA/B (hexamer 762 kDa) and oxoglutarate dehydrogenase complex (~2500 kDa).

### Generation of structural models for COX and AIFM1

Firstly, as no structure of bovine (dimeric) AIFM1 is currently available, a homology model was generated and structurally aligned based on the human dimeric AIFM1 structure (PDB: 4BUR) using Robetta. The final dimeric model of AIFM1 lacks the N-terminal region, containing residues 128–516 and 551–611 for both molecules. The N-terminal region of AIFM1 (res 55-124) was generated using trRosetta (Yang et al., 2020). Further, a monomeric COX structure was generated from the recently published bovine COX dimer (PDB: 1V54). The structure was modified by adding the missing NDUFA4 subunit, which was modelled and structurally aligned based on the human homolog (PDB: 5Z62 chain N) using Robetta. Likewise, missing residues (without transit peptides) for COX6B1 and COX5A subunits were modelled and added to the COX and added to the final COX structure used for docking.

### Cross-linking driven docking and analysis of a COX-AIFM12 complex

To generate a COX-AIFM1 structure, modified structures for COX, AIFM1 and the N-terminal domain of AIFM1 were used. Firstly, interaction interfaces and cross-links supporting a distinct complex formation were identified using DisVis (van Zundert and Bonvin, 2015). Active residues involved in an interface were computed additionally based on solvent accessible residue information. Solvent accessible residues (absolute and relative solvent accessibility ≥ 40%) were identified using the standalone program Naccess (© S. Hubbard and J. Thornton 1992-6). Structural docking with respective structures was done in Haddock (Karaca and Bonvin, 2011; van Zundert et al., 2016) using predicted active residues and cross-links as additional restraints. Different distance allowances were used based on the observed cross-link: DSSO = 35 Å, PhoX = 30 Å and DMTMM = 25 Å. The cluster supporting the cross-linking restraints best as well as with the best Haddock score was chosen as final model for a COX-AIFM1_2_ complex. Subsequently, the “Model Loops” of Chimera (Version 1.14rc) (Pettersen et al., 2004; Yang et al., 2012) was applied to model missing residues (125-127) and structurally connecting the N-terminal domain of AIFM1 (res 55-124) and respective AIFM1 protomer (28-516; 551-613). Interface residues of the resulting COX-AIFM1_2_ complex were identified using the Prodigy web service (Xue et al., 2016). To determine whether cytochrome *c* can still potentially bind to its COX binding site in the COX-AIFM1_2_ complex, this protein was structurally aligned based on a previously solved structure of cytochrome *c* docked to COX from bovine heart (Sato et al., 2016). Presented membrane boundaries for all presented structures were added using either the OPM (Lomize et al., 2012) webserver or MemprotPD (Newport et al., 2019).

#### Supplementary figures

**Figure S1.**
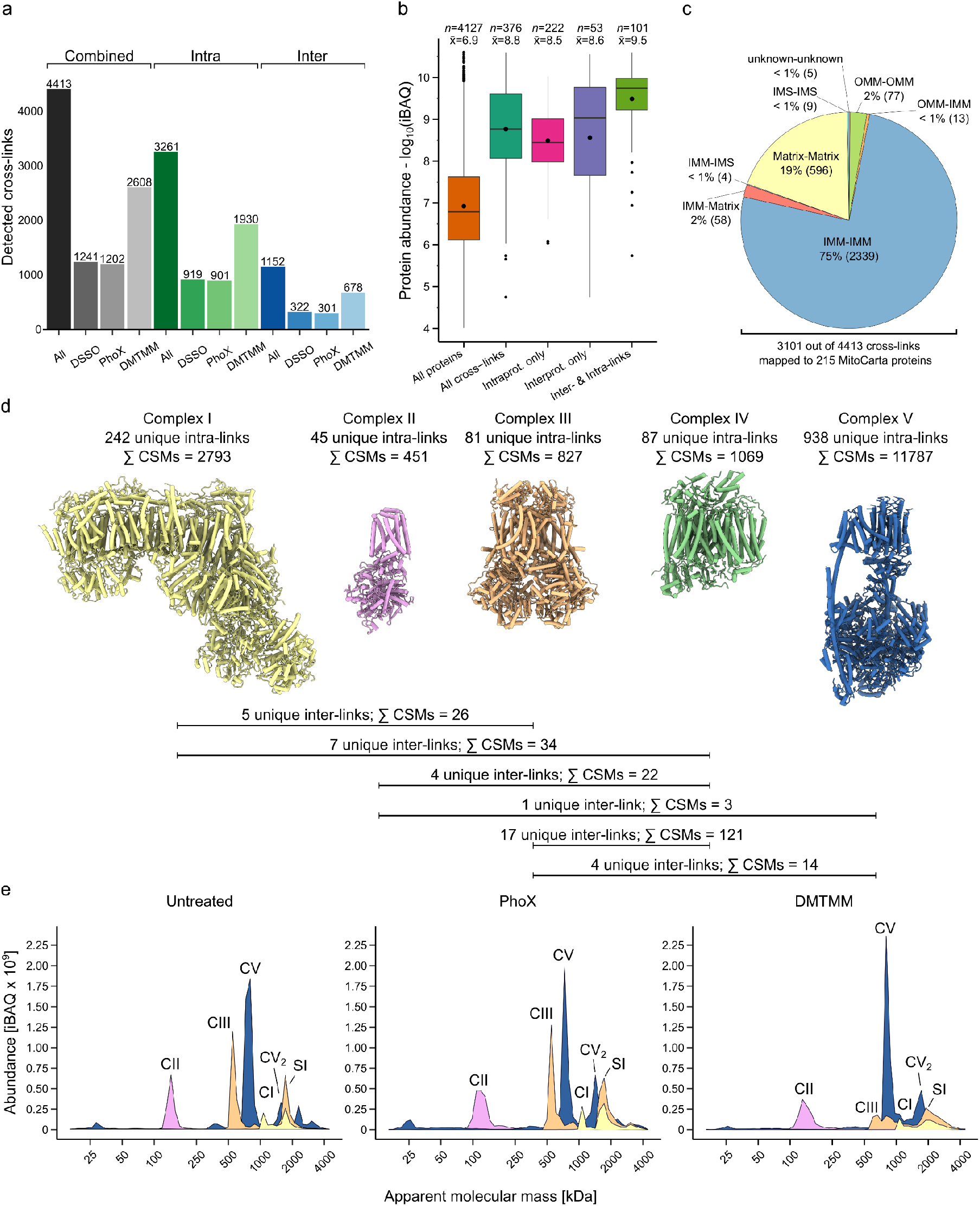
Overview of cross-linking and complexome profiling of bovine heart mitochondria (BHM). BHM were cross-linked in parallel with three different cross-linkers (DSSO, PhoX and DMTMM) and subjected to XL-MS analysis or complexome profiling. **a)** Overview of the number of unique cross-linked peptides identified with at least two CSMs, coming from protein-interactions that were identified with at least two cross-linkers. **b)** Observed cross-linked proteins are generally more abundant. Boxplot showing protein abundances for all identified proteins, all cross-linked proteins and proteins that have either only inter-, intra- or both (inter and intra) links. The number of proteins and the median iBAQ is indicated on top of each box. **c)** Pie chart showing the sub-mitochondrial localization of the cross-linked proteins identified based on their MitoCarta 3.0 annotation. **d)** Numbers of obtained cross-links for OXPHOS complexes I-V. Complexes are heavily cross-linked (~35 % of all detected cross-links). Cross-links are observed within subunits of the same complex (intra cross-links) but also between subunits of different OXPHOS complexes (inter-crosslinks). For structural representation, deposited structural models were chosen (PDB: 5LNK, 1ZOY, 1NTM, 1V54, 5ARA). **e)** Averaged migration profiles of the OXPHOS complexes CI, CII, CIII and CV without cross-linker treatment and after treatment with either the PhoX or DMTMM cross-linker using a 4-16% gradient BN gel. The profiles were obtained by plotting the relative abundance of the averaged subunits of each complex against the respective molecular mass. Peaks are annotated based on the molecular masses of CI, CII, CIII and CV. Upon addition of cross-linker, the OXPHOS complexes largely maintain their overall migration profile, and thus structural integrity.

**Figure S2.**
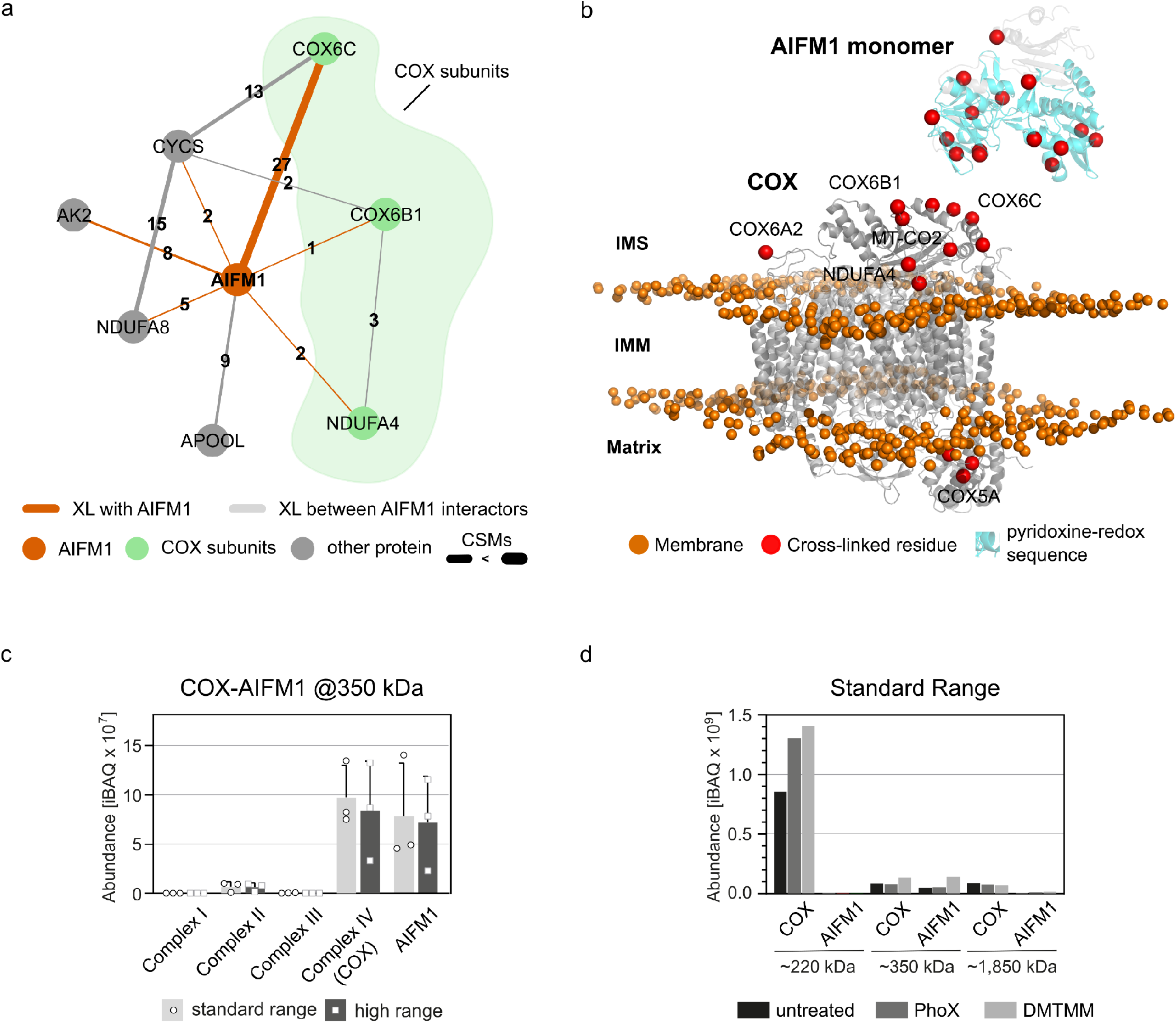
COX-AIFM1 interactions. **a)** The AIFM1 interactome as observed by XL-MS in intact mouse heart mitochondria adapted from (Liu et al., 2018). Numbers indicate the CSMs identification for each interaction. CSMs were summed from triplicates. **b)** Visualization of cross-linked residues (red spheres) of COX (gray) and truncated AIFM1 monomer (gray with residues of the Pyr-redox sequence being cyan) in our data on BHM. Orange spheres indicate the phosphate groups of a simulated lipid bilayer (IMM) which was structurally aligned based on the simulation for bovine monomeric COX (PDB: 6JY3) obtained from the MemProtMD server. **c)** Comparison of the amounts of respiratory chain complexes I to IV and AIFM1 at ~350 kDa representative of the COX-AIFM1_2_ complex. Complexome profiling was performed using a standard range BN-gel (4-16%) and a high range BN-gel (3-10%). The average of individual values±SD from all three conditions (untreated, PhoX and DMTMM cross-linked) is shown. **d)** Comparison of COX and AIFM1 abundances at ~220 kDa, ~350 kDa and ~1,850 kDa in complexome profiles using a standard BN-gel (4-16%) of BHM as prepared (untreated) and after cross-linking with PhoX and DMTMM. In **c** and **d,** iBAQ values of AIFM1 are divided by two to account for the dimer and the average of subunits of the respective complexes at the indicated approximate apparent masses in the migration profiles were taken as a measure for their abundance.

**Figure S3.**
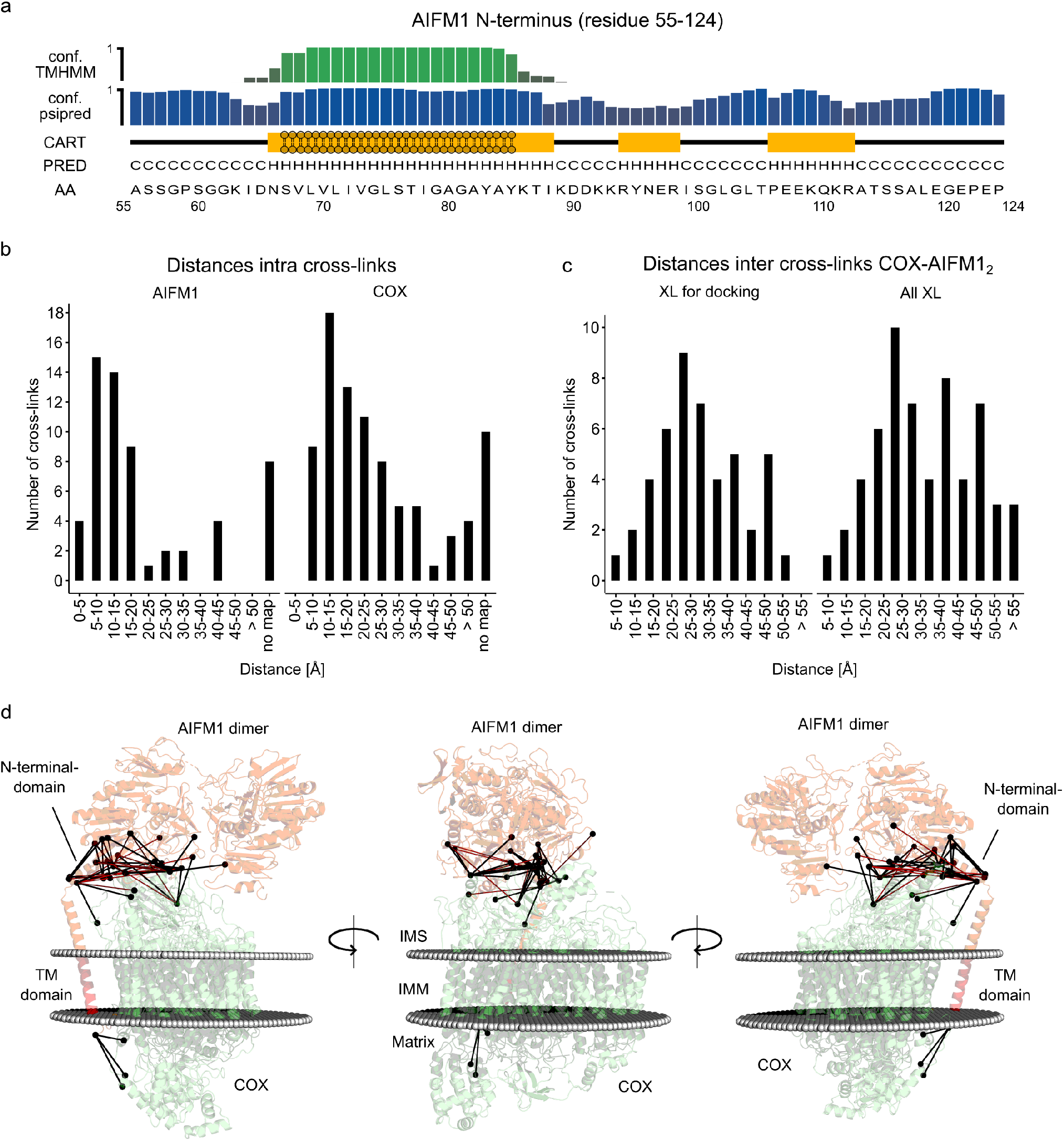
Structural properties of the N-terminal region of AIFM1 and cross-link distances in the COX-AIFM1 models. **a)** Secondary structure prediction of the N-terminal region of AIFM1 (res 55-124; sequence indicated under 4a). The upper bar plot shows the confidence of residues being trans-membrane residues (Score =1) or not within a membrane (Score < 0.6). The second bar plot shows the confidence of the secondary structure prediction (1 = highest, 0 = lowest) for each residue which is indicated under PRED as letter code (C = coil, H = helix) und visualized as cartoon under CART. **b)** Distance histogram of mapped cross-links (combination of DSSO, PhoX and DMTMM) for COX and AIFM1. AIFM1 includes distances for cross-links mapped on the AIFM1 dimer and the N-terminal region of AIFM1. For both structures, a number of cross-links are obtained within or attaching to missing regions of AIFM1 (N-terminus (55-127) and residue 517-550) and COX (N-terminus COX4l1). **c)** Cross-links (combination of DSSO, PhoX and DMTMM) used for structural docking and all observed cross-links for COX-AIFM1_2_ interaction were mapped onto the best complex model. For AIFM1, all obtained cross-links can be mapped onto the final structure of AIFM1, containing two AIFM1 protomers (res 128-516, 551-613) and one N-terminal region of one protomer (res 55-124). **d)** Visualization of cross-links between COX and AIFM1 which were used for structural docking of the COX-AIFM1_2_ complex. COX subunits are colored green, while AIFM1 protomers are colored orange. The TM-domain of the N-terminal domain of AIFM1 is highlighted in red. Boundaries of the IMM are indicated as gray spheres. Cross-links with a Cα-Cα distance ≤ 35 are colored in black, while cross-links with a mapped Cα-Cα distance ≥ 35 are colored in red. Cross-linked residues (lysine, glutamic acid or aspartic acid) are highlighted as black spheres.

**Figure S4.**
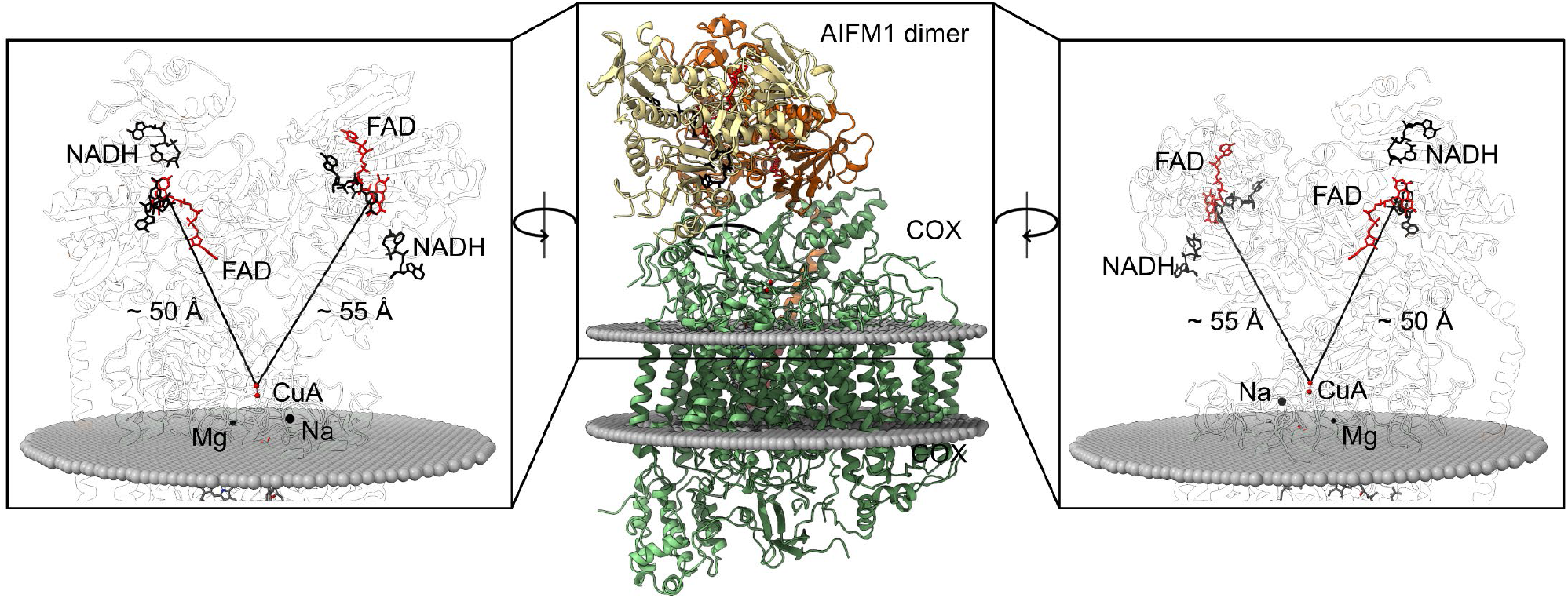
Mapping the distance from the isoalloxazine moiety of FAD in AIFM1 to the CuA center of COX. Cofactors were structurally aligned based on structures of AIFM1 (PDB: 4BUR) and COX (PDB: 1V54). AIFM1 protomers are orange and COX subunits are green (middle); red sticks represent FAD, black sticks represent NADH, Cu_A_, and Cu_B_ are red (left and right). Boundaries of the IMM are indicated as gray spheres.

